# Comparative evolutionary analysis and prediction of deleterious mutation patterns between sorghum and maize

**DOI:** 10.1101/777623

**Authors:** Roberto Lozano, Elodie Gazave, Jhonathan P.R. dos Santos, Markus Stetter, Ravi Valluru, Nonoy Bandillo, Samuel B. Fernandes, Patrick J. Brown, Nadia Shakoor, Todd C. Mockler, Jeffrey Ross-Ibarra, Edward S. Buckler, Michael A. Gore

## Abstract

Sorghum and maize share a close evolutionary history that can be explored through comparative genomics. To perform a large-scale comparison of the genomic variation between these two species, we analyzed 13 million variants identified from whole genome resequencing of 468 sorghum lines together with 25 million variants previously identified in 1,218 maize lines. Deleterious mutations in both species were prevalent in pericentromeric regions, enriched in non-syntenic genes, and present at low allele frequencies. A comparison of deleterious burden between sorghum and maize revealed that sorghum, in contrast to maize, departed from the “domestication cost” hypothesis that predicts a higher deleterious burden among domesticates compared to wild lines. Additionally, sorghum and maize population genetic summary statistics were used to predict a gene deleterious index with an accuracy higher than 0.5. This research represents a key step towards understanding the evolutionary dynamics of deleterious variants in sorghum and provides a comparative genomics framework to start prioritizing them for removal through genome editing and breeding.

## Main text

Sorghum (*Sorghum bicolor* L. Moench) and maize (*Zea mays* L.) are both members of the Poaceae family and often serve as a model system for comparative plant genomics. Their common Poaceae ancestor underwent a whole-genome duplication (WGD) event ~96 million years ago ^1^, and a second WGD in maize corresponds closely with its divergence 12 million years ago from sorghum ^2^. The role of polyploidization in maize diversification ^1^ makes the sorghum-maize system particularly powerful for comparative studies.

Archaeobotanical studies support a single sorghum domestication event around 3000 BC in Eastern Sudan ^3^, with genetic studies supporting a potential second independent domestication center in west Africa ^4,5^. Maize, in contrast, was domesticated once from teosinte in the Balsas river valley in central Mexico around 9000 years ago ^6,7^. While some orthologs between these two species experienced parallel selection during domestication ^8^, most domestication related genes appear to be drawn from a non-overlapping set ^9^. It is well established that maize experienced a decline in effective population size due to a domestication bottleneck ^10,11^ that increased the burden of deleterious alleles in the domesticate compared to teosinte ^12,13^. Evidence for reduced nucleotide diversity in landraces from a genetic bottleneck ^5^ or population size decline ^14^ has also been reported for sorghum.

Sorghum has a hermaphroditic inflorescence, contributing to its predominantly self-pollinating nature. Domesticated sorghum has an estimated outcrossing rate of only 7 to 20% ^9,15^, while the outcrossing rate tends to be higher (up to 70%^16^) for weedy and wild sorghum. In contrast, maize and its wild progenitor teosinte are monoecious and generally outcrossing at a rate of over 90%^17^. In this study, we performed a joint analysis of functional variation in sorghum and maize lines that span the wild-to-domesticated continuum to compare deleterious mutation accumulation in these two closely related species with distinct domestication histories and mating systems.

First, we conducted an extensive characterization of the levels and patterns of standing genetic variation in a diversity panel of 468 sorghum lines using whole-genome resequencing (WGS). These accessions represented a wide range of molecular and phenotypic diversity, including wild relatives, landraces, and improved breeding lines (Table S1). The average sequencing depth per line was 15× (Figure S1). Variants were called using the *Sorghum bicolor* reference genome (v3.1)^18^ (Figure S2), followed by filtering to obtain a high-quality core set of 13.2 million SNPs and 1.8 million indels (Figure S3).

Five morphological forms (races) have been previously defined within *S. bicolor*: bicolor, durra, guinea, caudatum and kafir. *S. bicolor* race bicolor was the first race to be domesticated ^19,20^, while the remaining four “modern” races exhibited parallel evolution for free-threshing grains. Additionally, these four races experienced a significant amount of introgression from wild relatives within various agro-climatic and geographical environments ^3,21,22^. Consistent with previous reports, we observed population structure at both the race (Fig 1A) and geographical levels using principal component analysis (PCA)^21,22^ (Fig S4). Linkage disequilibrium (LD) in the diversity panel decayed rapidly to 50% of its initial value within the first few Kb, reaching background levels (*r*^2^ = 0.1) around 300 kb (Fig 1B). We observed slight differences in LD decay rates (Fig S5A) and more striking allele frequency differentiation (F_ST_ 0.16-0.32; Fig S5B-C) among races, recapitulating sorghum’s domestication and adaptation history in variable environments. Local LD profiles revealed a significant increase in LD around the centromeres on all chromosomes except chromosome 1 (Fig1C), where a large genomic interval surrounding the centromere is missing from the reference genome ^23^.

**Figure 1.**
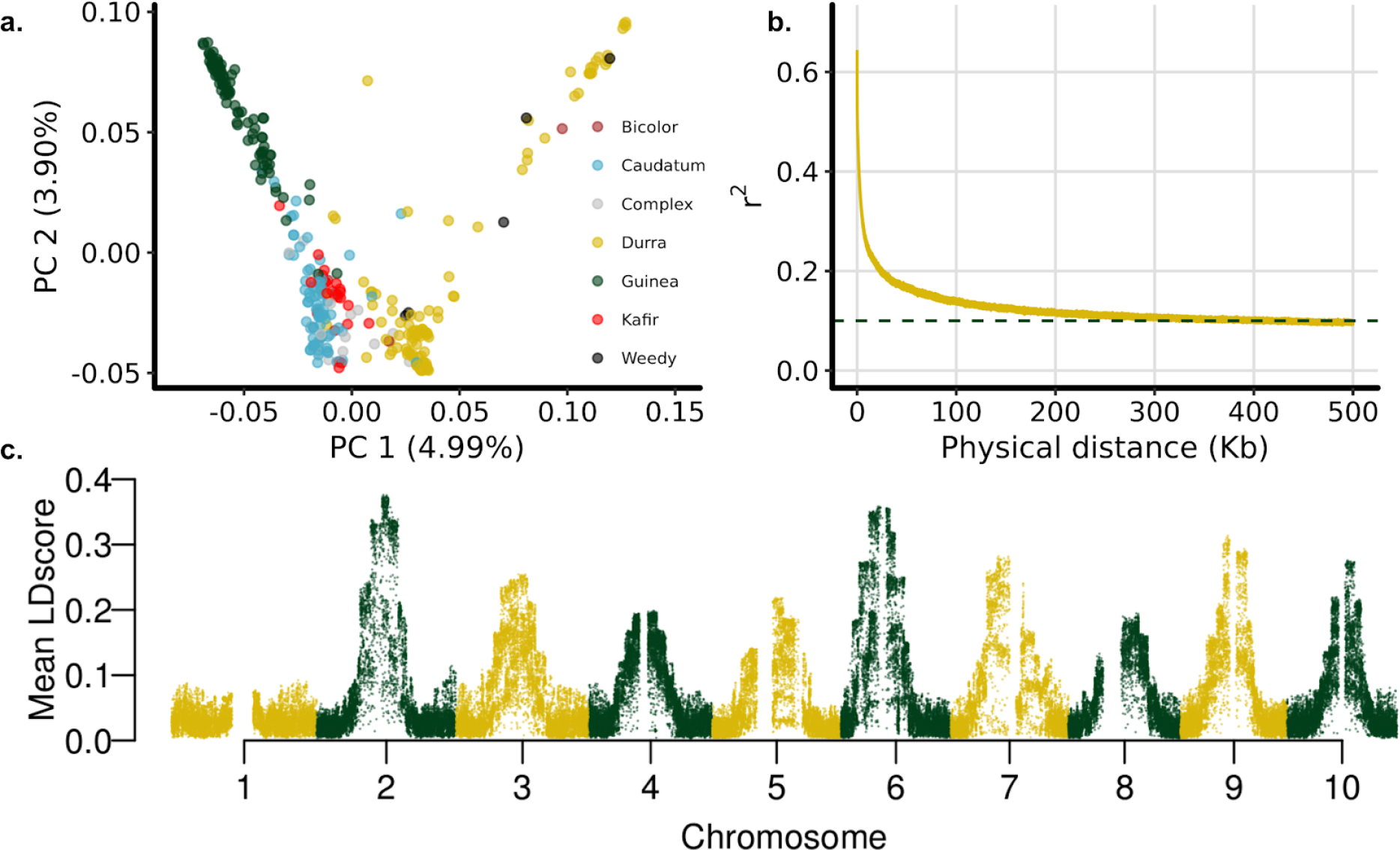
Population structure and linkage disequilibrium patterns in sorghum: **a**, Principal component analysis (PCA) of sorghum lines. **b**, LD decay rate as determined by the squared correlations of allele frequencies (*r*^*2*^) against physical distance (kb) between polymorphic SNP loci. **c**, Mean LD scores estimated with a 1Mb window. The reference genome is missing the majority of the centromeric region on chromosome 1.

To allow for a comparative analysis, genomic evolution and amino acid conservation modeling ^24,25^ was used to catalog candidate deleterious variants across both the sorghum and maize genomes. Genomic evolutionary rate profiling (GERP)^26^ identified 64.9 Mb ^25^ of the sorghum genome (9.49%) as evolutionarily constrained (GERP > 0). In maize, this value increased to 117 Mb^27^, but only represents 4.16% of its genome. As a complement, sorting intolerant from tolerant (SIFT)^28^ scores were calculated to predict the effect of amino acid substitutions on protein function for both maize and sorghum. In sorghum, of the 459,188 SNPs identified within exons, 19% (87,684) were considered putatively deleterious (SIFT < 0.05). Similarly, 8% (53,583) of the exonic SNPs in maize were annotated as deleterious. As expected, we found that a large percentage of the variants inside coding regions were also evolutionarily constrained (GERP > 0) in both sorghum (44%) and maize (53%) (Figure S6).

We split coding variants into five categories based on SIFT and GERP scores: deleterious (GERP >= 2, SIFT <0.05), non-conserved deleterious (GERP <2, SIFT <0.05), stop mutations (either gain or loss), tolerated (nonsynonymous, SIFT > 0.05), and synonymous mutations (Figure S7). Similar to maize^29^, the sorghum derived allele frequency (DAF) spectrum (Figure S8) showed that the categories of deleterious mutations exhibited an excess of low frequency variants compared to non-deleterious variants. As in maize ^27^, deleterious mutations were also enriched in sorghum pericentromeric regions where suppressed recombination makes it difficult for the organism to purge these variants (Figure S9).

After maize and sorghum split from a common ancestor, the former went through a whole genome duplication around ~12 million years ago ^30^. Once both maize subgenomes were combined together within a single nucleus, the process of “fractionation” commenced, whereby one copy of each duplicated gene pair tends to be lost ^30^. We investigated whether the fractionation and syntenic status between these two species is associated with the accumulation of deleterious variants. We found the proportion of variants within each of the five categories to be similar between fractionated and non-fractionated genes in both maize and sorghum, but the number of deleterious variants was significantly higher in non-syntenic genes for both species (Figure 2A). Our data suggest that non-syntenic genes, potentially less essential, carry deleterious variants at higher levels in both species.

**Figure 2.**
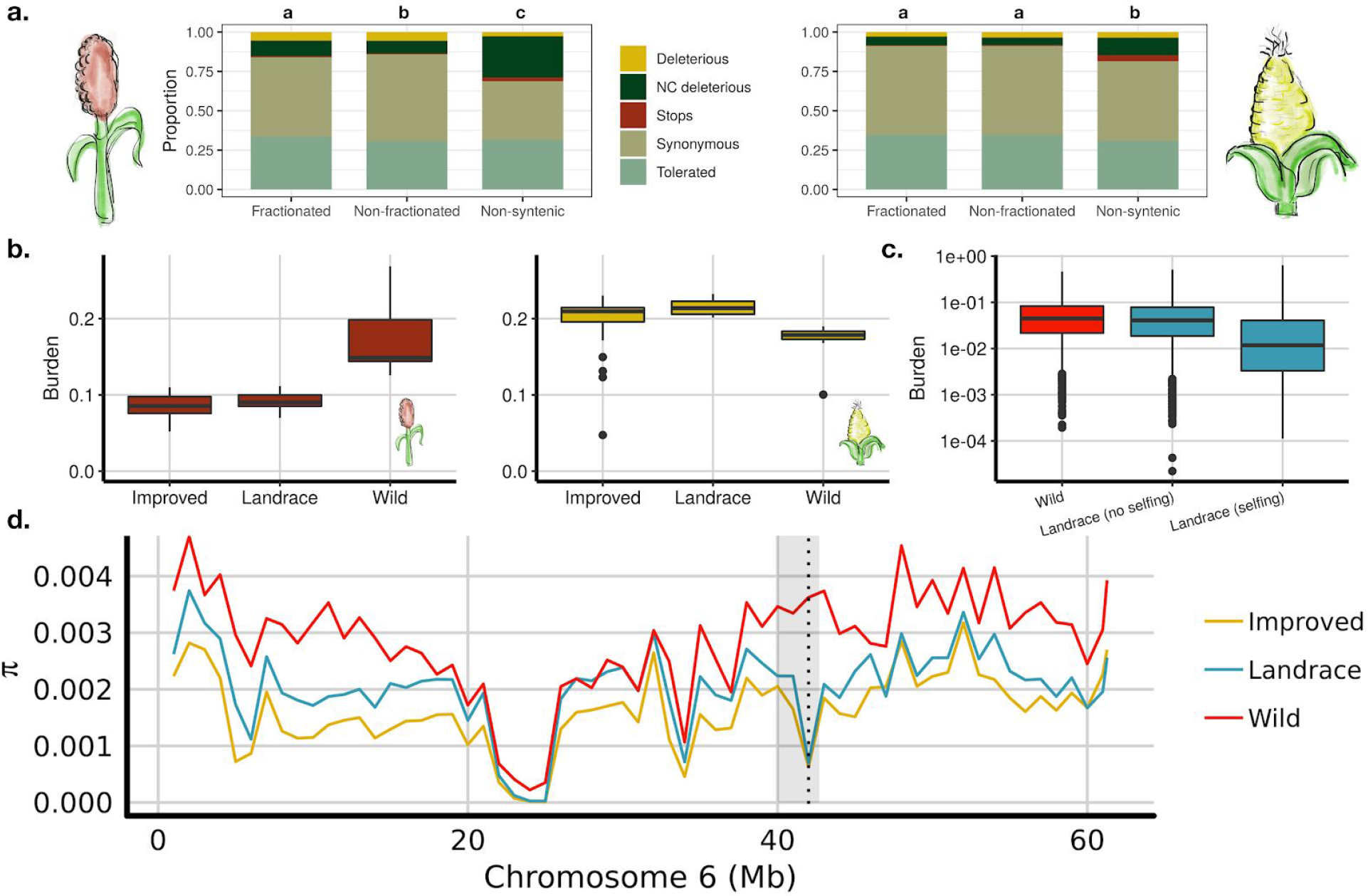
Comparative analysis of deleterious alleles in sorghum and maize: **a**, Distribution of variants inside coding sequences for sorghum (left) and maize (right). Variants were divided into five categories: synonymous mutations (mutations that do not change the encoded amino acid), tolerated mutations (nonsynonymous mutations, SIFT > 0.05), stop-codon mutations (either gain or loss), non-conserved deleterious mutations (NC, deleterious, SIFT <0.05, GERP < 2), and conserved deleterious mutations (Deleterious, SIFT <0.05, GERP > 2). The genes were divided into three categories: fractionated (sorghum gene with one syntelog in maize), non-fractionated (sorghum gene with two syntelogs in maize), and non-syntenic. Different letters represent significant differences between groups using a two-proportion Z-test. **b**, Deleterious burden was calculated for sorghum (left) and maize (right). Burden was calculated as the sum of homozygous derived alleles at deleterious sites over the total number of sites available for each individual (recessive model). **c**, Distribution of genetic load under a simulation scenario. Simulations were based on the mean population size 10,000 years ago for sorghum landrace and wild lines inferred by SMC++^36^. The effect of increased selfing during sorghum domestication was explored by changing the inbreeding probability from 0 (no selfing) to 1 (selfing). **d**, Nucleotide diversity (*π*) estimates across chromosome 6 were calculated in 1Mb bins for improved (yellow), landrace (blue), and wild lines (red). The dotted line marks a major decline in nucleotide diversity for improved and landrace lines relative to wild lines. This genomic region (42Mb) colocalizes with *dw2* and *ma1*, which are known height and maturity loci ^21^. See Figure S10 for plots of all 10 chromosomes.

We also explored the deleterious burden of wild relatives, landraces, and improved lines in both maize and sorghum. The “cost of domestication” hypothesis ^31^ predicts that the process of domestication and crop improvement is likely to result in an increased number of deleterious variants in the genome. Our results in maize agreed with previous findings of an excess of overall deleterious alleles among improved maize lines relative to the wild relative *Z. mays* ssp. *parviglumis* ^*12*^ (Figure 2B). In sorghum, however, wild relatives had the highest accumulation of deleterious alleles. When analyzing each sorghum race independently, we observed three tiers of deleterious mutations, whereby wild relatives had the highest burden followed by the bicolor race and weedy lines. The modern agro-climatic adapted races showed the lowest burden (Figure S10). This departure could be explained partially by the inherent difference in mating systems of sorghum (selfing) and maize (outcrossing), especially from the transition to higher selfing rates in sorghum post-domestication. Notably, simulations that include an increase in the rate of selfing after domestication recapitulated the decrease in genetic load of landraces compared to wild taxa, which coheres with our interpretation of the data (Figure 2C–S11). This difference in burden is not driven by a lower genetic diversity observed in wild sorghum, as they follow the expected pattern (wild > landrace > improved) (Figure 2D–S12). Additionally, Hamblin *et al* ^32^ showed that sorghum domestication is a complex process that might have included ancestral population structure, multiple domestication events, and/or introgressions from wild relatives. A recent study ^14^ has further shown that genetic load could have been reduced in modern lines by introgression with wild sorghum relatives. We hypothesize that the differences observed in load accumulation between maize and sorghum might be driven by both their different mating systems and domestication histories.

Given the recent development of supervised machine learning applied to population genetics and genomics inference ^33,34^, we evaluated the efficacy of convolutional neural networks (CNNs) in building an evolutionary model capable of predicting a deleterious index (average SIFT score) and syntenic state for sorghum genes. We split the sorghum genome into segments with each gene as a centroid and calculated 12 features per window, four of which were derived from maize. We incorporated predictors of functional importance, conservation of synteny and fractionation of maize genes, levels of gene expression variance, and several molecular evolution statistics (Figure 3).

**Figure 3.**
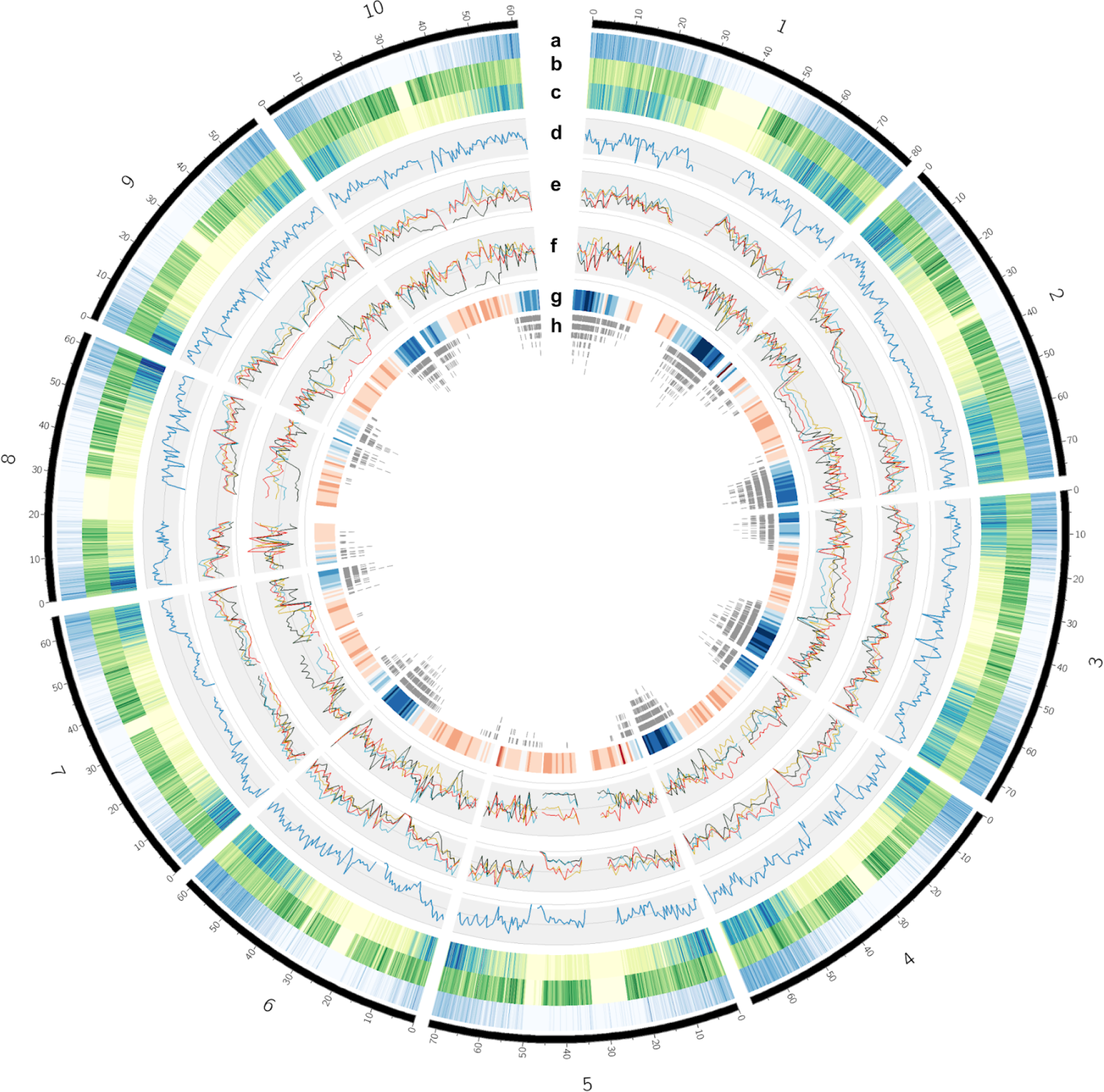
Genomic landscape of sorghum. **a**, Gene density heatmap. **b**, SNP density. **c**, Indel density. **d**, rho (ρ) population recombination rates were calculated using FASTEPRR in 500kb windows. **e**, Nucleotide diversity (π) in 1Mb windows for each sorghum race, the color codes are the same as used in Figure 1A, (red=kafir, blue=caudatum, yellow=durra, dark green=guinea). **f**, Tajima’s D for each sorghum race. **g**, Average GERP score per 1Mb window, with a divergent red-blue scale where blue represents high GERP score. **h**, Genomic distribution of sorghum non-fractionated genes.

Through the implementation of this model, we predicted the average SIFT score per gene, a statistic that reflects how likely a genic mutation would be deleterious. Our model had a prediction accuracy of 0.53 and outperformed linear regression models by 10% (Figure S13). The importance of individual features was assessed with a “leave one variable out” approach. We found that four features were most impactful. Two of them—average GERP score and number of variants in the CDS—were expected to be important as they both reflect the strength of purifying selection at a locus (Figure 4A–B). The other two impactful features were RNA expression variance in sorghum and maize syntelogs, supporting previous research in maize where the dysregulation of expression was correlated with rare-allele burden^35^. We also used the model to predict the syntenic state of the focal gene in each window with an AUC ~0.9. Two maize related features were most relevant: *ssw*, a measure of conservation between two genomes using short kmer alignments (see methods), and maize nucleotide diversity (π) (Figure 4C).

**Figure 4.**
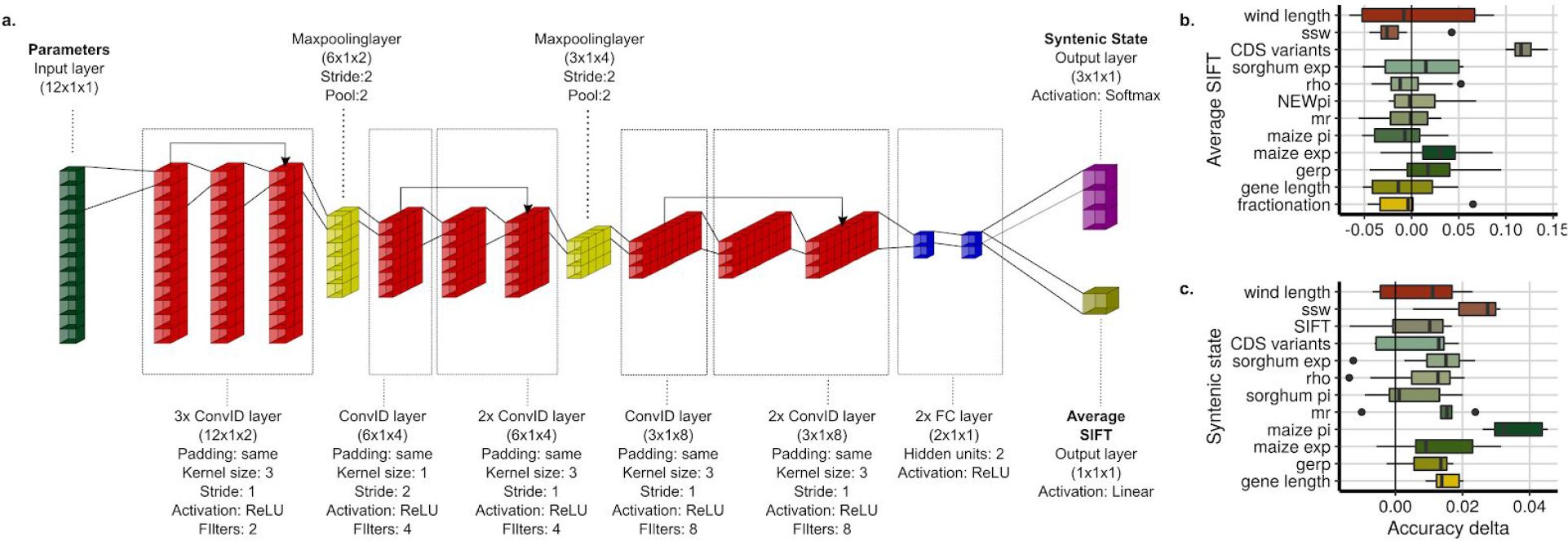
Convolutional neural network architecture. **a**, The sorghum genome was divided into ~34k windows, with each gene serving as a centroid. For each window, 12 features were calculated. Four of these features were calculated to exploit the syntenic relationships between maize and sorghum; nucleotide diversity (π) of the syntenic segment in maize (maize pi), ssw short alignments (*ssw*; see methods) between sorghum and maize, coefficient of variation (CV) of the expression from syntenic maize genes (maize exp), and syntenic state (fractionation). The remaining features were calculated in sorghum: nucleotide diversity (sorghum pi), conservation score (mean GERP score per window, gerp), neutral substitution rate (nr) per focal gene, average SIFT score (SIFT), number of variants in the CDS of the focal gene (CDS variants), population recombination rates (ρ), coefficient of variation (CV) of the expression of the focal gene across different developmental stages and treatments (sorghum exp), window length (wind length), and focal gene length (gene length). The CNN was trained to predict the average SIFT score per gene and the syntenic state of the focal gene (fractionated, non-fractionated, or non-syntenic). To assess the relative importance of each feature, we calculated the decrease in accuracy (Accuracy delta) after removing a single feature for predicting **b**, average SIFT score (continuous) and **c**, syntenic state (categorical).

In summary, we performed a joint variant analysis of sorghum and maize genomic variation across the wild-to-domesticated continuum to highlight their differences in genetic load accumulation. We also prototyped an evolutionary model using supervised machine learning that exploited the syntenic relationship between these two species. Overall, we constructed a landscape of the sorghum genome that can be used to support GWAS results, variance component estimates, and inform whole-genome prediction in a comparative genomics framework.

## Methods

### Plant material

The plant material used in this study is comprised of sorghum lines that represent a wide range of molecular and phenotypic diversity. Overall, 485 lines (468 of which are unique) were used to build a sorghum haplotype map. Only unique lines were considered for further analysis. The material has three different origins, Mace et al. ^5^ (*n* = 42), TERRA-MEPP project (*n* = 240, https://terra-mepp.illinois.edu) and TERRA-REF (*n* = 203, http://terraref.org/). Information for each line is included in Table S1.

### Sequencing and variant identification

The samples from all three datasets were sequenced using Illumina paired-end reads of 100bp or 150bp in length. The sequencing depth for each line ranged from 0.03 to 74.15× (Table S1, Figure S1). This was estimated based on the number of reads, expected sorghum genome size (732Mb), and read length. Variant calling was performed using Sentieon’s DNAseq pipeline 201711.03 ^37^, a proprietary implementation of the GATK variant calling pipeline ^38^. Briefly, BWA ^39^ version 0.7.13 was used to align the raw sequence reads to the BT×623 reference genome v3.1 available in Phytozome ^18^. Next, duplicated reads were removed and local realignment was performed around the indels. Then, base quality score recalibration (BQSR) was also performed using SNP markers identified in a previous sorghum hapmap ^25^ built with the 240 TERRA-MEPP accessions. Finally, gvcf files were generated using the Haplotyper in the emit_mode gvcf. The 485 gvcfs were then jointly called to produce a single VCF file using the GVCFtyper mode. For more details on the pipeline see Figure S2A and code in the GitHub repository (https://github.com/GoreLab/Sorghum-HapMap).

After variant calling, a vcf file containing ~41 million markers was generated. Of these, 35,025,902 were biallelic SNPs and 3,486,787 were biallelic indels (See Figure S2B for details). Hard filters were applied using GATK best practices recommendations. Briefly, SNP markers having a QualByDepth (QD) < 2, FisherStrand (FS) > 60.0, RMSMappingQuality (MQ) < 40.0, MappingQualityRankSumTest (MQRankSum) < −12.5, or ReadPosRankSumTest (ReadPosRankSum) < −8.0 were removed. Similarly, indels with QualByDepth (QD) < 2.0, FisherStrand (FS) > 200.0, or ReadPosRankSumTest (ReadPosRankSum) < −20.0 were also removed (Figure S2B). The average inbreeding coefficient (F) at filtering STAGE 2 was 0.68. This value contrasted with our expectation of high homozygosity for sorghum inbred lines. We applied to the data an “allele balance” filter (see GitHub repository vcfaddanot.py) that calculated the ratio between the number of reads supporting the heterozygous (het) calls and the total number of reads:

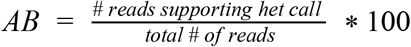

According to previous research^40^ most of the variants with an AB value above 30 could be validated using Sanger sequencing. Applying this same threshold, heterozygous calls with a value of AB lower than 30 were masked. Additionally, vcftools ^41^ was used to mask genotypes with a read depth (minDP) lower than 4, markers that were monomorphic, and markers with call rates lower than 50%. Finally, SNPs with an inbreeding coefficient lower than 0 were removed. The inbreeding coefficient per marker was calculated as:

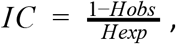

where H_obs_ and H_exp_ are the observed and expected heterozygosity under Hardy-Weinberg Equilibrium (HWE). Finally, the missing genotypes were imputed and phased into haplotypes using Beagle 4.1 ^42^ using a default window size of 50,000 SNPs and an N_e_ = 150,000. In total, 13,170,712 SNPs and 1,804,397 indels were called and imputed for the 485 accessions. Variants were called for all lines, however, only unique lines with sequencing depth higher than 2X were considered for further analysis (*n* = 466).

### Linkage Disequilibrium

Linkage disequilibrium (LD) decay was calculated using PopLDdecay v3.31 (https://github.com/BGI-shenzhen/PopLDdecay). Measures of LD (*r*^*2*^) were calculated for the entire population, but also for each chromosome and subpopulation. Pairwise *r*^*2*^ estimates were calculated from the unimputed SNP dataset with MAF > 0.05 and maximum missing rate < 0.25. LDscores were calculated using the Genome-wide Complex Trait Analysis (GCTA) suite ^43,44^ with default settings.

### Deleterious mutations

#### Candidate deleterious alleles identification

We used sorting intolerant from tolerant (SIFT) ^45^ to annotate sorghum and maize SNP data. The SIFT algorithm predicts whether an amino acid substitution is deleterious to protein function ^28^. SIFT uses protein alignments to identify conserved amino acids and provide a score of the putative deleterious effect for each position of the protein. These scores range from 0 to 1, and positions with a SIFT score < 0.05 are predicted to be deleterious.

In sorghum, GERP (genomic evolutionary rate profiling) scores ^26^ estimated from the alignment of six species including *Zea mays*, *Oryza sativa*, *Setaria italica*, *Brachypodium distachyon*, *Hordeum vulgare* and *Musa acuminate* ^46^ were used to identify evolutionary constrained nucleotides. From the ~13 million SNPs, 455,546 variants were located within coding sequence (CDS) and were split into 5 categories based on SIFT and GERP scores as follows: synonymous mutations (mutations that do not change the encoded amino acid), tolerated mutations (nonsynonymous mutations, SIFT > 0.05), stop-codon mutations (either gain or loss), non-conserved deleterious mutations (SIFT <0.05, GERP < 2), and conserved deleterious mutations (SIFT <0.05, GERP > 2).

The derived site frequency spectrum was calculated for the 455,546 CDS variants using both *Zea mays* (AGPv3.22 annotation) ^47^ and *Setaria italica (Setaria italica v2.2, DOE-JGI,* http://phytozome.jgi.doe.gov/) as outgroups to determine the ancestral state. Briefly, each sorghum gene was aligned with its syntenic orthologs ^48^ using clustal omega ^49^ in both maize and setaria. For each variant, the corresponding nucleotide in maize (both subgenomes) and setaria were identified and the sts-ufs software ^50^ was used to infer the probability of the derived vs. ancestral allelic state. The same approach was used to annotate deleterious alleles in maize. Briefly, maize HapMap v3 ^51^ (1200 individuals) was used to annotate deleterious alleles. Only markers with the “LLD” flag present and the “NI5” flag absent were used, as suggested by the authors. The “LLD” flag includes SNPs that are confirmed to be in LD with GBS anchor markers and the “NI5” flag is used to mark SNPs within 5bp of an indel. Together, the “LLD” flag present and the “NI5” flag absent represents the cleanest marker set ^51^. Maize annotation on the v3 reference genome (AGPv3.22) ^47^ was used to calculate SIFT scores. GERP scores were available from Rodgers-Melnick et al. ^52^. Sorghum and setaria were used as outgroups for maize to infer the derived/ancestral allele using the sts-ufs software^50^.

Utilizing a single reference genome for a species when calculating GERP and SIFT scores introduces a bias that underestimates the number of deleterious alleles in domesticated material versus wild relatives ^12^. It has been previously shown that annotation of markers for which the reference allele is derived might be unreliable ^53^. To mitigate this bias, we calculated the probability that each reference-derived allele would have been classified as it was (deleterious, NC deleterious, etc) had the reference allele been ancestral separately for maize and sorghum. Briefly, all the alleles for which BTx623 (sorghum reference genome genotype) or B73 (maize reference genome genotype) where ancestral were divided into bins of 1% derived allele frequency and the fraction of reference-ancestral sites in each functional category was calculated. Those fractions were used as weights for all the reference-derived sites.

#### Deleterious Burden calculations

The fitness for a variant was calculated relative to the ancestral state ^54^. The ancestral allele was set as the non-deleterious marker (coded as 0 in the dosage matrix) and the derived allele as the potential deleterious allele. We then calculated the burden of each line using the unimputed marker matrix. The homozygous burden (recessive model) was defined as the sum of homozygous deleterious sites (coded as 2 in the dosage file) over the total deleterious sites called for each individual (not considering missing genotype calls). The heterozygous burden was defined as the sum of heterozygous deleterious sites (coded as 1 in the dosage file) over the total deleterious sites called for each individual. The individuals used for the comparison between improved lines, landraces, and wild relatives for maize and sorghum are included in Table S2.

#### Deleterious Burden simulations

To understand the potential reason for decreased burden in landraces compared to wild sorghum, we simulated different scenarios. To parameterize our simulations to fit the empirical data, we estimated the population history of sorghum using SMC++ ^36^. We selected five lines from each group (wild, landrace, and improved) to ensure equal sampling. We used the repeat masking of the sorghum reference genome version 3.0.1 to create input files for each line of a group as a distinct lineage. We used forward in time simulations in fwdpy11 (https://github.com/molpopgen/fwdpy11), a Python package using the fwdpp library ^55^ to simulate a diploid random mating population. We simulated a 100 Mb region where deleterious mutations occurred at 1% of a neutral mutation rate of 3×10^−8^ mutations/bp/generation and a recombination rate equal to the neutral mutation rate.

The effect size of deleterious mutations followed an exponential distribution with a mean of −0.05. Based on the mean population size 10,000 years ago for the landrace and wild lines inferred by SMC++, we simulated an equilibrium population having a constant size of N_anc_= 11270 for 10 N_anc_ generations before changing parameters. The demographies used followed the inferred demography for the last 10,000 years. We tested the effect of increased inbreeding from selfing during sorghum domestication by changing the inbreeding probability from 0 (completely outcrossing) to 1 (complete inbreeding) and from 0.5 to 1. We replicated each parameter combination 100 times. At the end of the simulation, we recoded the fitness of 50 random individuals. We calculated burden as the difference in fitness of an individual and the fittest individual with the same ancestral inbreeding.

### Convolutional neural network model

The sorghum genome was divided into windows containing a single focal gene. Then, several summary statistics from both sorghum and maize were calculated for each window. These windows were randomly split into training, validation, and test sets and a CNN framework was used to build a model for predicting average deleterious score (SIFT) and syntenic state. Syntenic state refers to whether the focal sorghum gene in each window is fractionated, non-fractionated, or non-syntenic when compared with maize. Each of the features used in the model are detailed in the next section.

#### Defining sorghum genome windows

The midpoint distance between adjacent genes was calculated, allowing for the calculation of a series of intervals that covered each chromosome from start to end. In total, the sorghum genome was divided into 34,028 windows (Table S3).

#### Nucleotide diversity (pi), Tajima’s D and per site Fst

Nucleotide diversity ^56^ and Tajimas’s D were calculated with the sorghum and maize SNPs using vcftools --site-pi ^41^S, --TajimaD, and --weir-fst-pop. Using bedtools, we calculated the average pi/TD/Fst value per window counting every nucleotide site inside the window and giving a value of 0 for monomorphic positions.

#### GERP (Genomic evolutionary rate profiling)

Sorghum GERP ^26^ scores were estimated from the alignment of six species including *Zea mays*, *Oryza sativa*, *Setaria italica*, *Brachypodium distachyon*, *Hordeum vulgare* and *Musa acuminate* ^46^. For further information, please refer to Valluru et al. ^25^. We used maize GERP scores calculated previously by Rodgers-Melnick et al. ^52^. Average GERP scores were calculated per window. Additionally, average neutral substitution rates (nr, sum of the tree branch lengths) were calculated for each focal gene.

#### SIFT (Sorting Intolerant From Tolerant)

Sorghum SIFT annotations were calculated with a sorghum database created using the Sbicolor_313.v3.1 gene annotation. Only primary transcripts were taken into consideration. Sift4g was used to evaluate the 13 million sorghum variants. Raw SIFT results are available through the CyVerse repository associated with this manuscript (http://datacommons.cyverse.org/browse/iplant/home/shared/GoreLab/dataFromPubs). Maize SIFT scores were calculated on a subset of the HapMap v3 markers ^51^, only including those with the “LLD” flag present and the “NI5” flag absent (29 million variants). For maize, the pre-computed SIFT database on the v3 reference genome (AGPv3.22) was used. Raw SIFT results are available in the CyVerse repository associated with this manuscript. Average SIFT scores per gene were calculated.

#### SSW (SSW alignment between sorghum and maize)

The sorghum reference genome was aligned to the maize genome with a sliding window of 20bp using the Complete Striped Smith Waterman library (SSW; https://github.com/mengyao/Complete-Striped-Smith-Waterman-Library) for faster alignments. We reported the number of times each 20bp tag from sorghum aligned to maize, the maximum alignment score of the 20bp tag, and the average of multiple alignment scores of each 20bp tag. For each window used in the evolutionary model, the average maximum alignment score was calculated and used as a proxy of local conservation between the sorghum and maize genomes.

#### Rho (ρ, population recombination rate)

Population recombination rates were calculated in the entire panel using the software FASTEPRR^57^. and also to calculate the recombination rate for each of the gene windows used to build the evolutionary model (FASTEPRR_segments.R).

#### Sorghum and maize gene expression

Six sorghum (PRE0023, PRE0028, PRE0146, PRE0295, PRE1441 and Grass1) and two (B73 and Mo17) maize lines were grown under the same greenhouse experimental conditions. Plant material for RNA sequencing was collected from two tissues (growing point and leaves) at four developmental stages (three, five, six, and seven leaf stage) during day and night conditions. In total, 192 and 64 sample combinations of genotypes, developmental stages, tissue types, and diurnal conditions for sorghum and maize were obtained, respectively. Samples were processed according to Kremling et al. ^35^. Briefly, RNA was extracted using TRIzol (Invitrogen) with Direct-zol columns (Zymo Research) and 3′ RNA-seq libraries were prepared from 500 ng total RNA in 96-well plates on an NXp liquid handler (Beckman Coulter) using QuantSeq FWD kits (Lexogen) according to the manufacturer’s instructions. Libraries were pooled to 96-plex and were sequenced with 90 bp single-end reads using Illumina TruSeq primers on an Illumina NextSeq 500 with v2 chemistry at the Cornell University Biotechnology Resource Center (BRC).

The first 12 bp and Illumina Truseq adapter remnants were removed from each read using Trimmomatic version 0.32. The splice-aware STAR aligner v.2.4.2a was used to align reads against either the sorghum v3.1 or the maize AGPv3 reference genome annotations, allowing reads to map to at most 10 locations (-outFilterMultimapNmax 10) with at most 4% mismatches (–outFilterMismatchNoverLmax 0.04), while filtering out all non-canonical intron motifs (–outFilterIntronMotifs RemoveNoncanonicalUnannotated). Default settings from STAR v.2.4.2a aligner were used to obtain gene-level counts (--quantModel GeneCounts) from the resulting BAM files.

#### CNN architecture

A deep residual convolutional neural network was developed (DeepEvolution), tailored to predict different population genetics parameters in classification and regression problems. The DeepEvolution architecture is composed of one input layer, nine one-dimensional convolutional layers, two max pooling layers, two fully connected layers, and one output layer (Figure 4a). In order to better exploit deeper low and high level representations from the input space, residual blocks over the network were used as motivated by the ResNet architecture ^58^ and batch normalization after convolutional layers. ReLU activations were applied on the hidden layers to better leverage nonlinearities from the population genetics parameters space. A decreasing number of channels and filter sizes were chosen, which is common in many efficient computer vision algorithms like the AlexNet architecture ^59^. For the regression problem of predicting average SIFT score, a linear activation function was used in the output layer. In the classification framework, the softmax activation function was implemented to predict the probability of the three classes of syntenic state (2 copies, non-fractionated; 1 copy, fractionated; 0 copy, non-syntenic). The network for average SIFT score was optimized using the Adam algorithm with a learning rate of 0.001 to minimize the mean-squared error function for regression, while for and syntenic state categorical cross-entropy was used for classification. Additional details about the model architecture, cross-validation, and feature importance analysis can be reviewed in the documented code (https://github.com/GoreLab/Sorghum-HapMap/tree/master/CNN/codes). The DeepEvolution model was fitted using the library keras 2.1.6 and python 3.6. To compare with accuracy obtained from the CNN, we used linear regression (average SIFT score) and multinomial logistic regression (syntenic state) models using the same predictors as in the CNN.

## Data availability

The raw sequencing data for the TERRA-MEPP lines are available through the NCBI BioProject PRJNA513297. Mace et al. raw data are available through the BioProject PRJNA182489, TERRA-REF raw data are available through the data commons database at CyVerse: http://datacommons.cyverse.org/browse/iplant/home/shared/terraref. Gene expression raw data are available through the Bioproject PRJNA503076. SIFT raw results and VCF files among others are available through the CyVerse repository: http://datacommons.cyverse.org/browse/iplant/home/shared/GoreLab/dataFromPubs (DOI:10.####/pending once accepted). Code used throughout the article is available at the GitHub repository: https://github.com/GoreLab/Sorghum-HapMap.

## Acknowledgments

Thank you to James Schnable for the maize, sorghum and setaria ortholog lists. The authors would also like to thank the Ross-Ibarra lab at UC Davis for helpful comments and sound advice on an earlier draft of this manuscript. The information, data, or work presented herein was funded in part by the Advanced Research Projects Agency-Energy (ARPA-E), U.S. Department of Energy, under Award Numbers DE-AR0000598, DE-AR0000661 and DE-AR0000594. The views and opinions of authors expressed herein do not necessarily state or reflect those of the United States Government or any agency thereof. This work was carried out with the support of Cooperative Research Program for Agriculture Science and Technology Development (Project No. PJ01321305) Rural Development Administration, Republic of Korea. For JPRDS, this work was partially supported by FAPESP grants 2017/03625-2 and 2017/25674-5 / CAPES (São Paulo Research Foundation) Finance Code 001 / Conselho Nacional de Desenvolvimento Científico e Tecnológico (CNPq).

