## Supplementary_figures for "Comparative evolutionary analysis and prediction of deleterious mutation patterns between sorghum and maize"

Ravi Valluru

Present Address: International Maize and Wheat Improvement Center (CIMMYT), El Batan 56237, México

Elodie Gazave

Present Address: Institute of Biotechnology, Cornell University, Ithaca, NY 14853, USA

Nonoy Bandillo

Present Address: Department of Plant Sciences, North Dakota State University, Fargo, ND 58105, USA

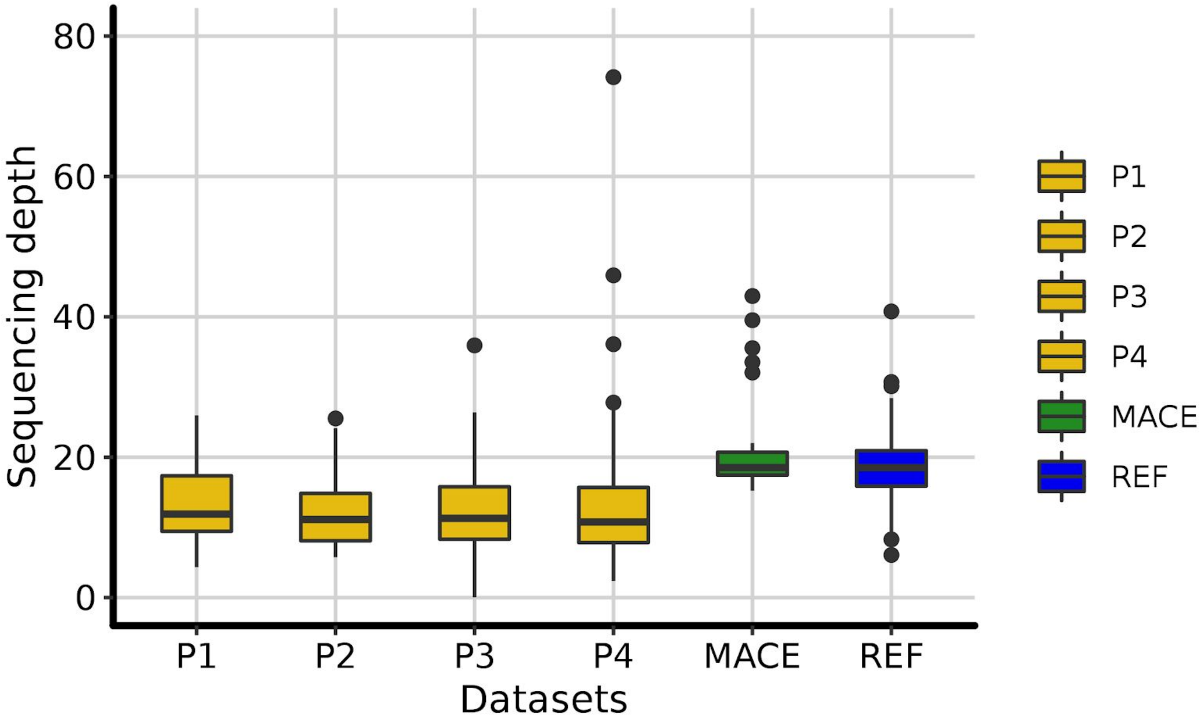


**Figure S1. Sequencing depth across sorghum datasets.** Sequencing depth was calculated as the number of reads multiplied by the number of bases per read, then divided by the estimated size of the sorghum genome (~732Mb). The different batches are Mace et al. (MACE) [1], TERRA-REF (REF), and the four batches from the TERRA-MEPP project (P1-P4). See Table S1 for more details.

**(A)**

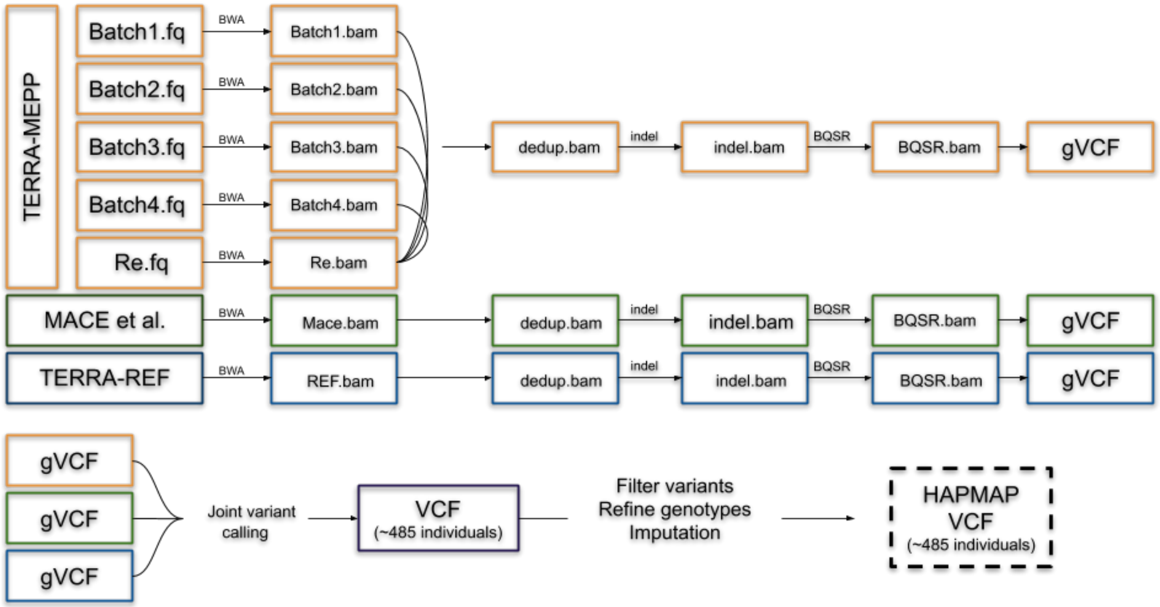


**(B)**

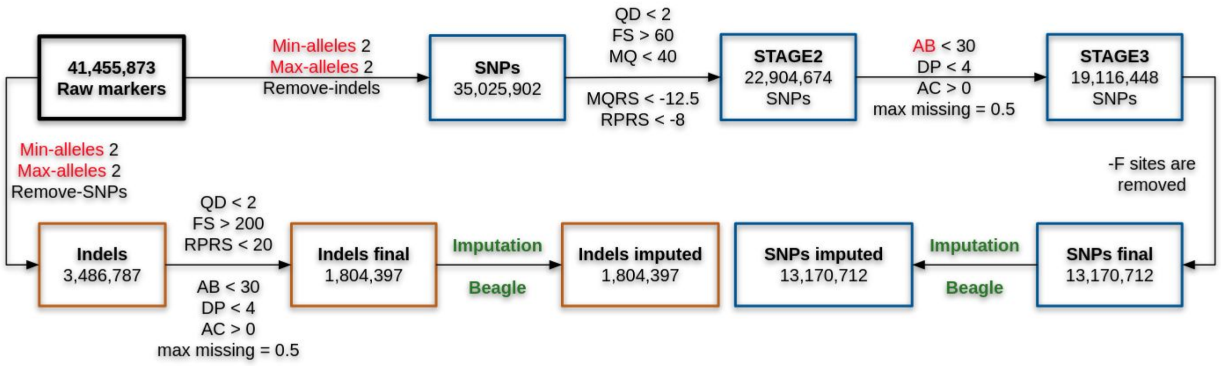


**Figure S2. Variant calling and filtering pipeline. (A)** Overview of the different steps taken to call variants on the 485 sorghum lines (includes duplicates). Each dataset has a different color, Mace et al. (green), TERRA-REF (blue), and TERRA-MEPP (orange). The TERRA-MEPP samples were split into 4 different batches and each batch was sequenced twice to increase the sequencing depth. **(B)** The variant filtering pipeline was used to filter the original 41 million variants down to the final set of 13M SNPs and 1.8M indels. Variant annotation was performed on the SNPs Final (unimputed) dataset. The details of each filter can be found in the methods section and VCF files for SNPs and Indels are available at: http://datacommons.cyverse.org/browse/iplant/home/shared/GoreLab/dataFromPubs/Lozano_MaizeSorghum_2019.

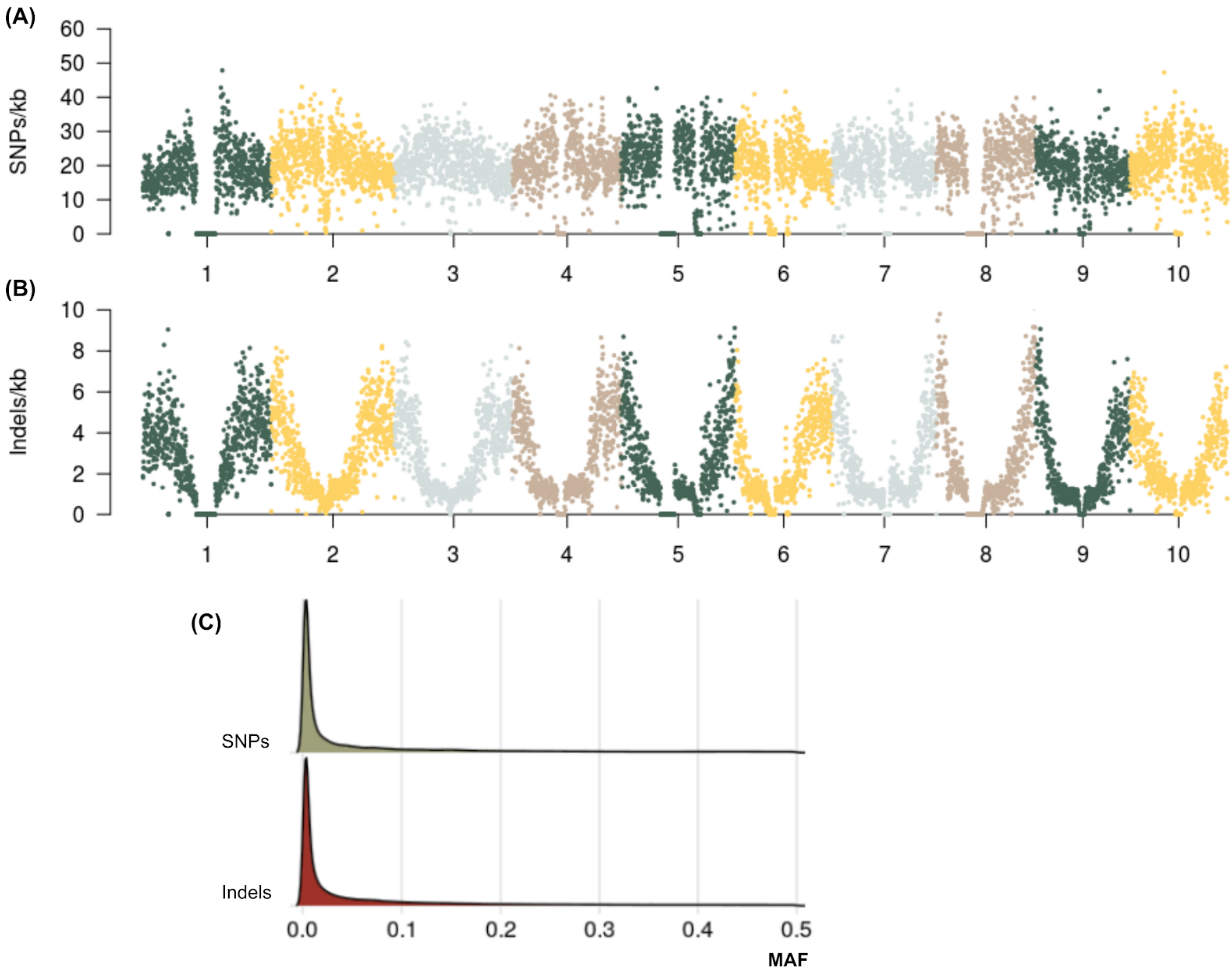


**Figure S3. SNP and indel distribution.** (**A**) SNP and indels **(B)** density for each of the 10 sorghum chromosomes. The genome was divided into 1Mb bins and each point represents the number of markers per kb in each bin. **(C)** Minor allele frequency (MAF) distribution of SNPs and indels.

**(A)**

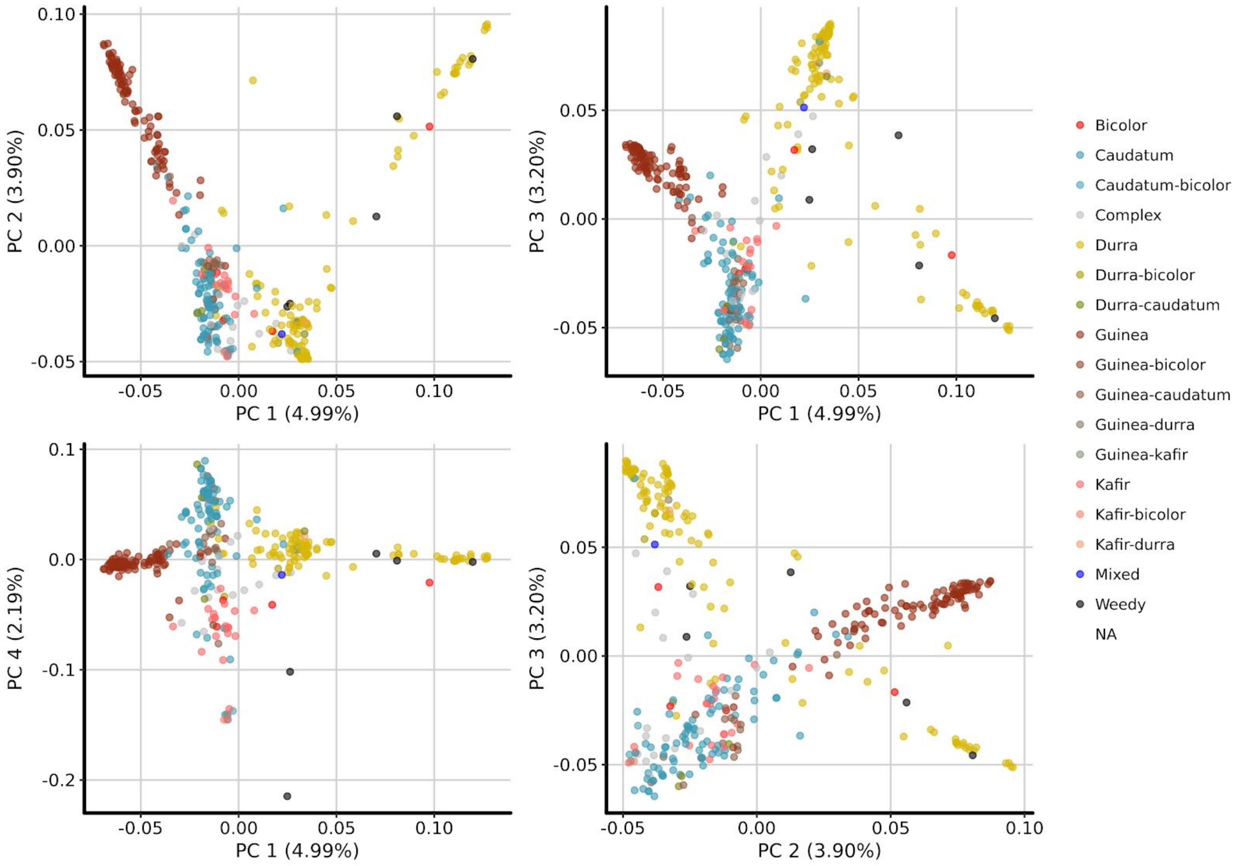
PC 1 (4.99%)
PC 2 (3.90%)
PC 3 (3.20%)
PC 4 (2.19%)
Bicolor
Caudatum
Caudatum-bicolor
Complex
Durra
Durra-bicolor
Durra-caudatum
Guinea
Guinea-bicolor
Guinea-caudatum
Guinea-durra
Guinea-kafir
Kafir
Kafir-bicolor
Kafir-durra
Mixed
Weedy
NA

**(B)**

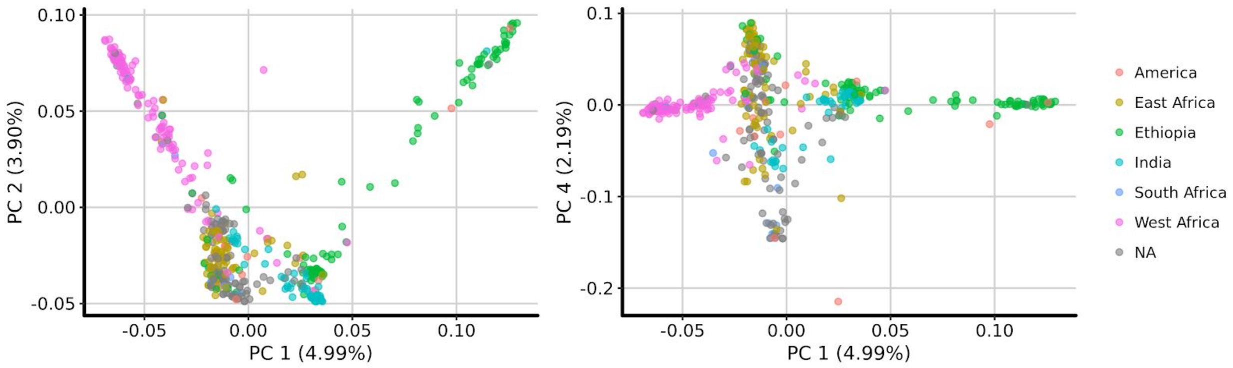
PC 1 (4.99%)
PC 2 (3.90%)
PC 4 (2.19%)
America
East Africa
Ethiopia
India
South Africa
West Africa
NA

**(C)**

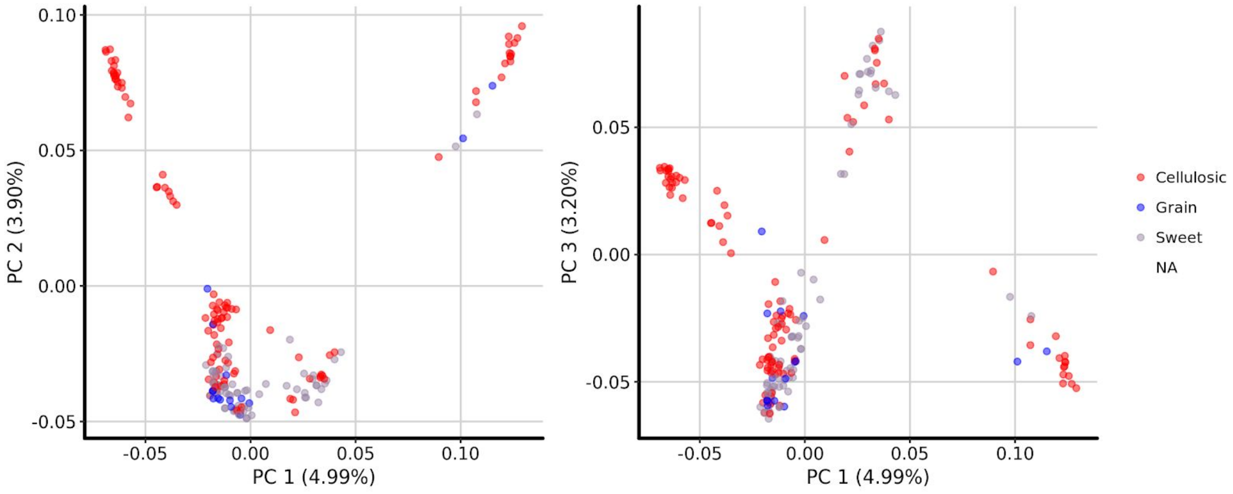


**(D)**

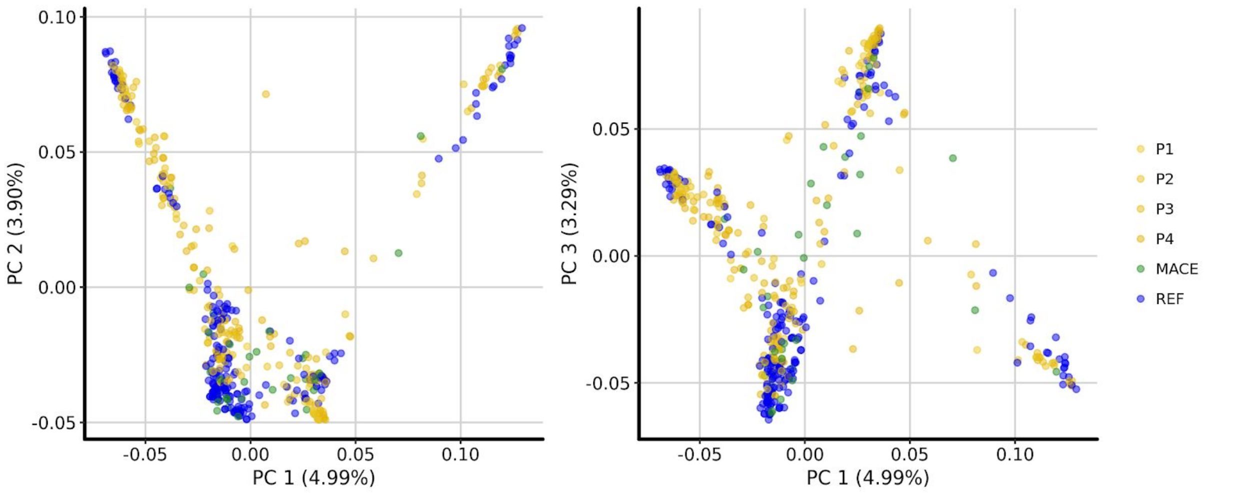


**Figure S4. Principal component analysis (PCA) on the SNP genotype matrix. (A)** PCA colored by race, the first 4 principal components (PCs) are shown **(B)** Geographical separation recapitulates race differences. **(C)** No major genetic separation was observed between sweet, cellulosic, and grain sorghum. **(D)** No evident batch effects were detected using the first 3 PCs. The different colors are samples from Mace et al. (MACE) [1], TERRA-REF (REF), and the four batches from the TERRA-MEPP project (P1-P4).

**(A)**

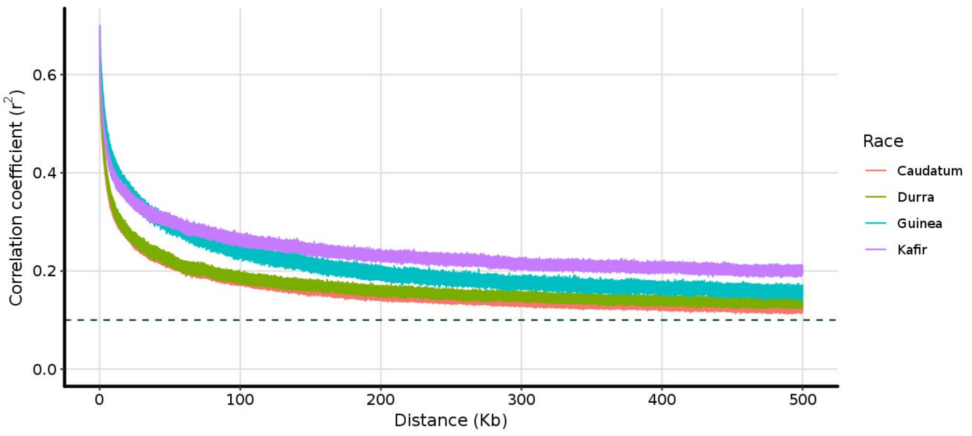


**(B)**

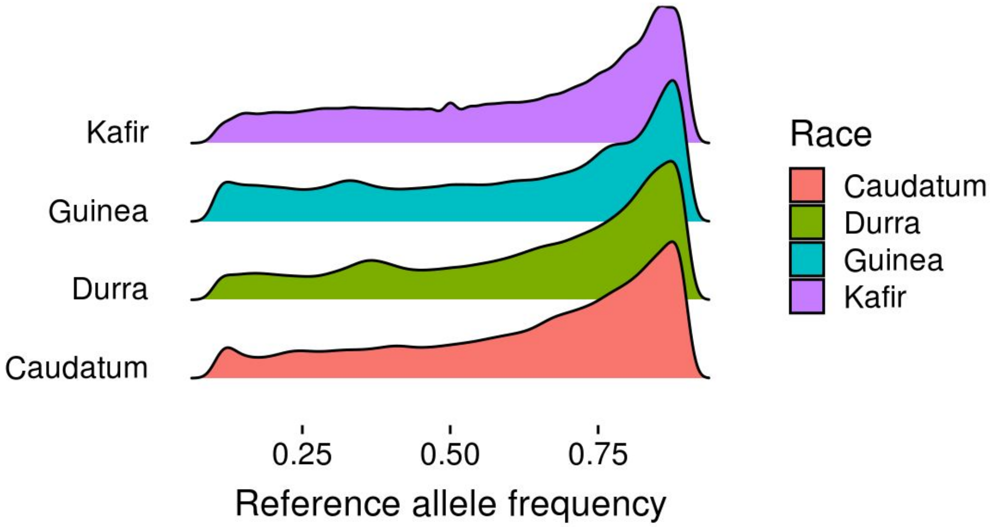


**(C)**

| **Fst** | **Caudatum** | **Durra** | **Guinea** | **Kafir** |
|---|---|---|---|---|
| **Caudatum** | | **-** | **-** | **-** |
| **Durra** | 0.189 | | **-** | **-** |
| **Guinea** | 0.238 | 0.308 | | **-** |
| **Kafir** | 0.159 | 0.246 | 0.318 | |

**(D)**

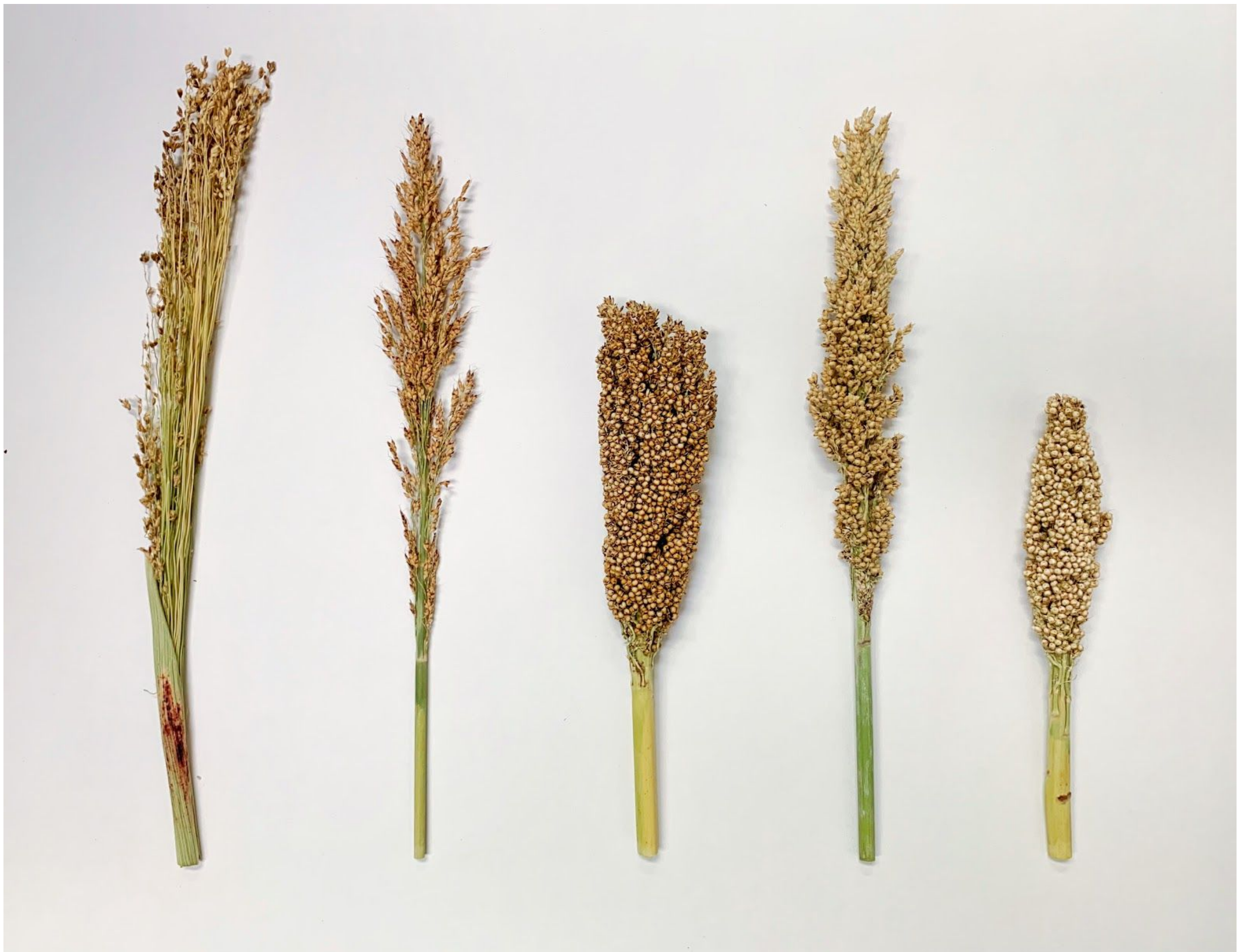

**Figure S5. Population genetics statistics of sorghum races**. **(A)** Linkage disequilibrium (LD) decay was calculated using PopLDdecay (https://github.com/BGI-shenzhen/PopLDdecay) for each of the four races: Caudatum (*n*=79), Durra (*n*=111), Guinea (*n*=93), and Kafir (*n*=23). **(B)** Reference allele frequency distribution of common alleles (minor allele frequency > 0.10) for each race. **(C)** Weighted pairwise fixation index ($F_{st}$) calculated using SNPrelate[2] on a random subset of 100,000 genome-wide SNP markers. **(D)** Image showing variation in panicle architecture of the different sorghum races. From left to right: Bicolor, Guinea, Caudatum, Kafir and Durra. Photo by Nadia Shakoor.

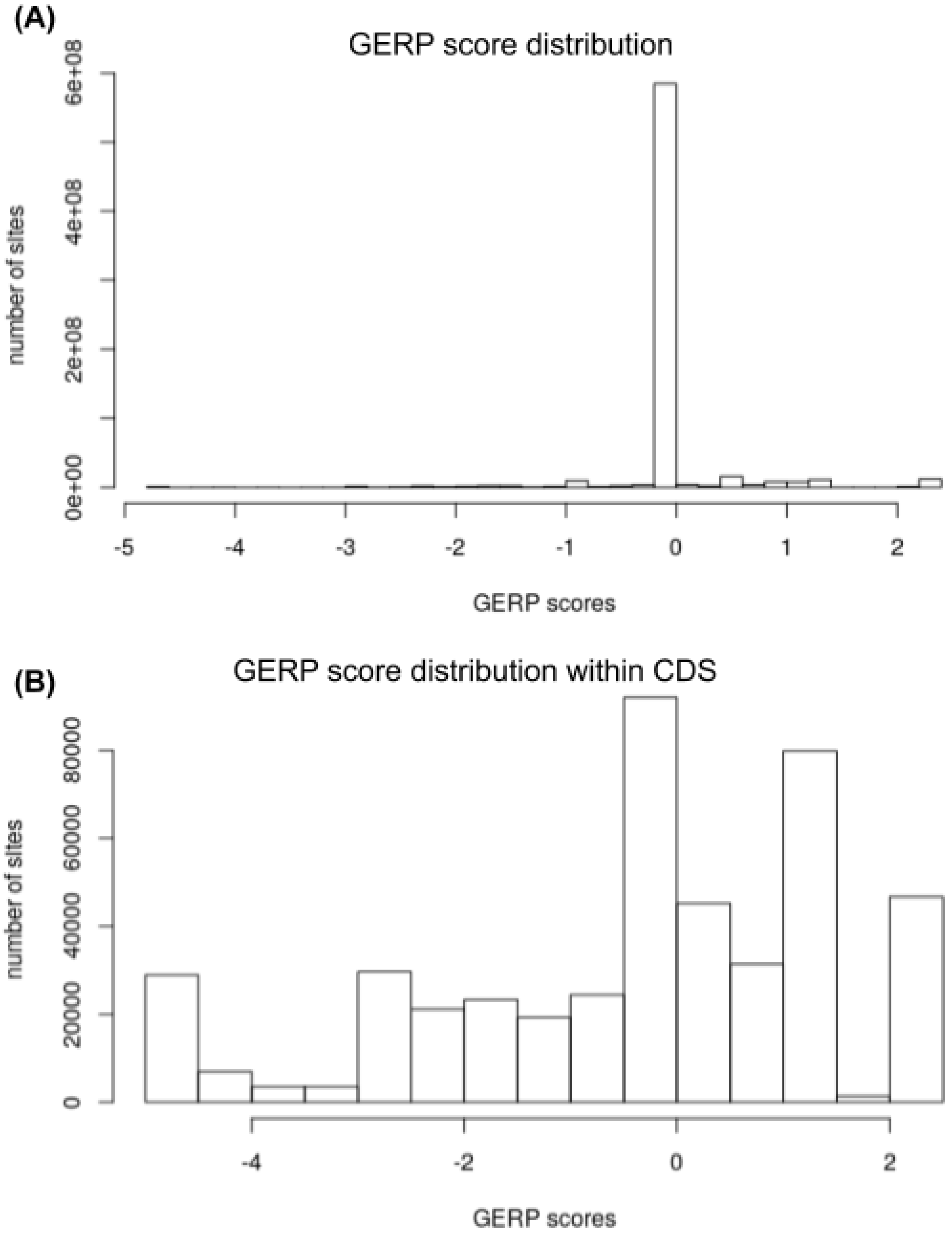


**Figure S6. GERP score distribution in the sorghum genome**. (A) GERP (genomic evolutionary rate profiling) score distribution across the genome. (B) GERP score distribution within coding sequence (CDS).

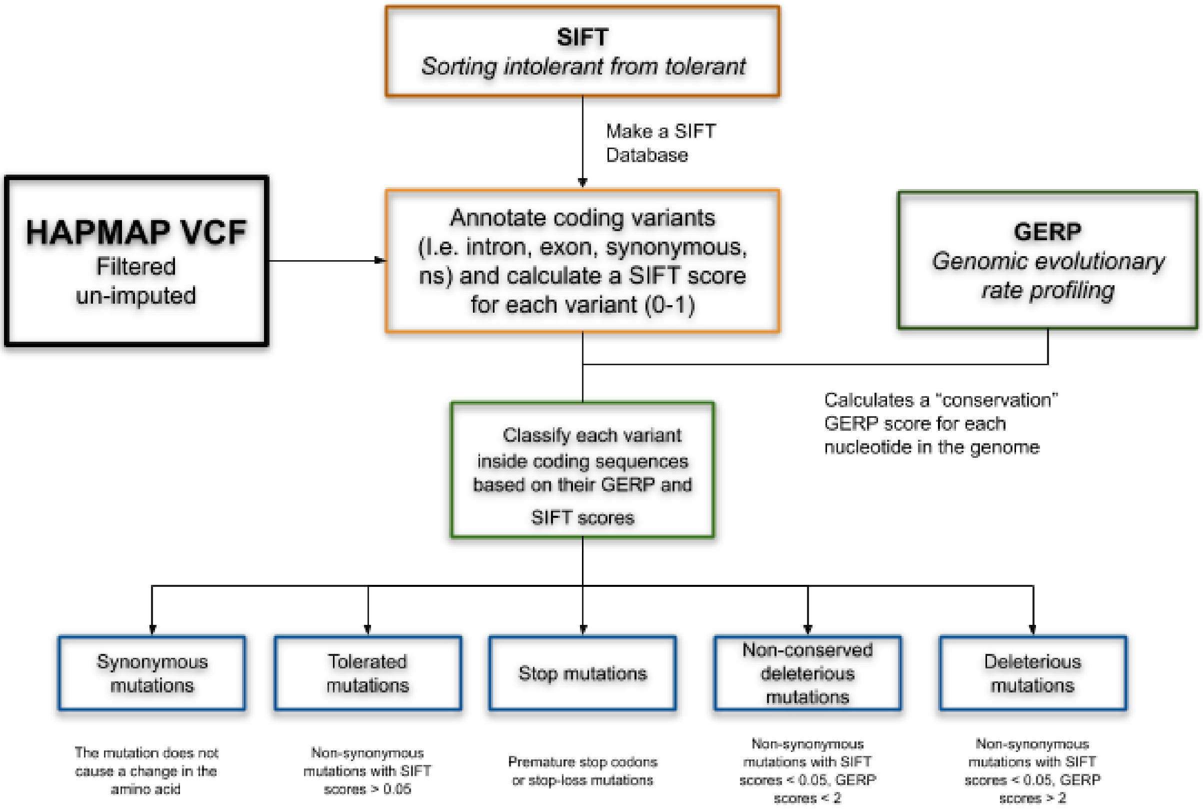


**Figure S7. Deleterious mutation annotation pipeline.** Workflow of the deleterious mutation annotation pipeline and the five categories into which coding variants were separated. Synonymous mutations (mutations that do not change the encoded amino acid), tolerated mutations (nonsynonymous mutations, SIFT > 0.05), stop-codon mutations (either gain or loss), non-conserved deleterious mutations (SIFT <0.05, GERP < 2), and conserved deleterious mutations (SIFT <0.05, GERP > 2).

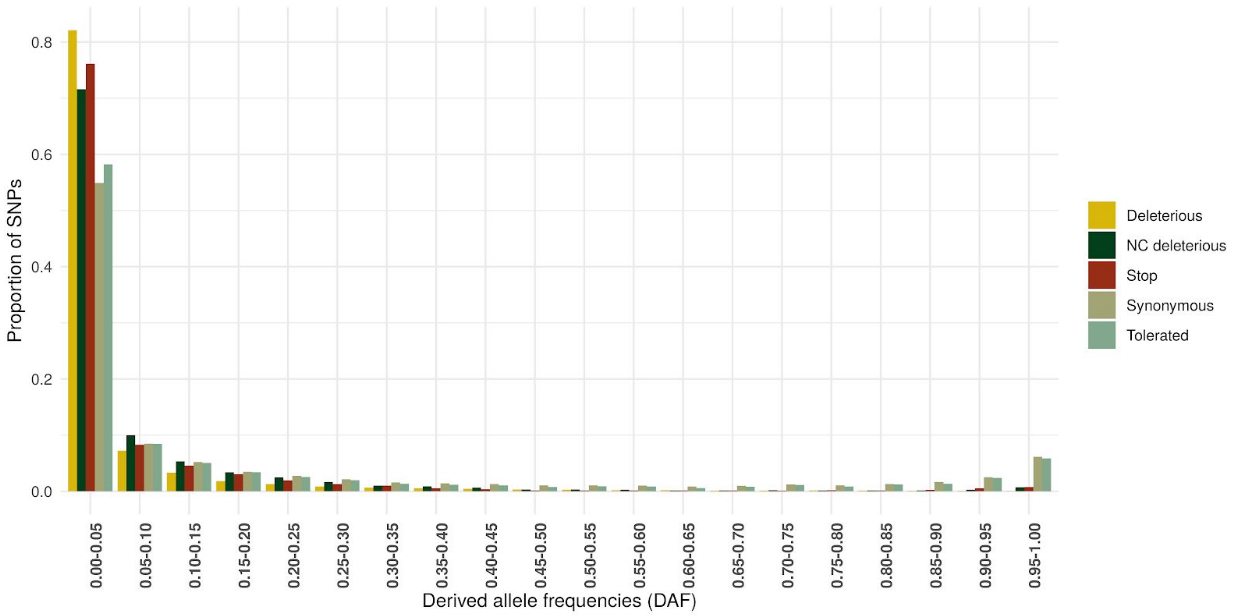


**Figure S8. Derived site frequency spectrum of sorghum coding variants.** Variants were divided in five categories: synonymous mutations (mutations that do not change the encoded amino acid), tolerated mutations (nonsynonymous mutations, SIFT > 0.05), stop-codon mutations (either gain or loss), non-conserved deleterious mutations (NC, deleterious, SIFT <0.05, GERP < 2), and conserved deleterious mutations (Deleterious, SIFT <0.05, GERP > 2). Derived/ancestral alleles were identified using a maximum likelihood method (sts-ufs[3]) under a Kimura two-parameter model using maize and *Setaria italica* as outgroups.

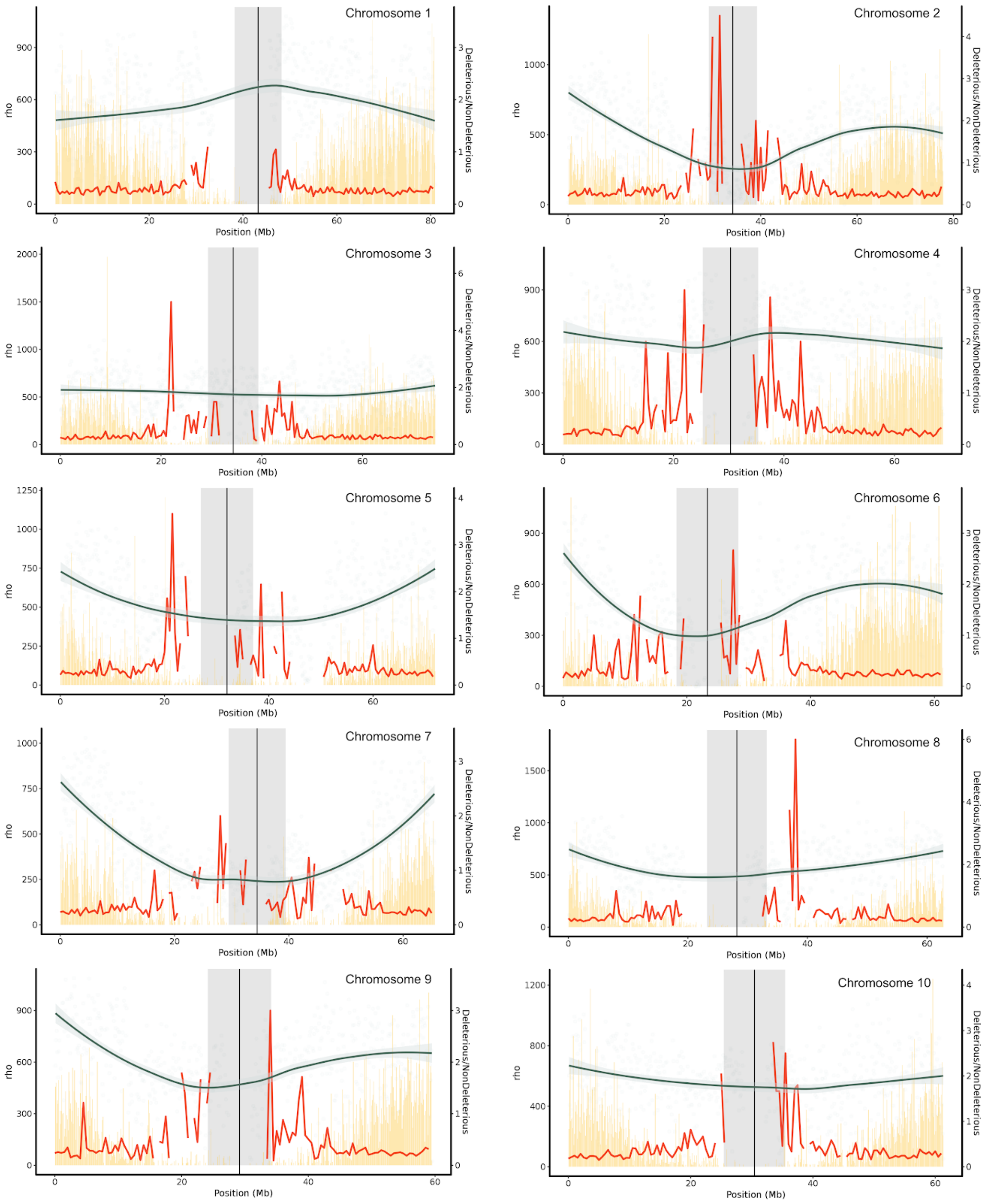


**Figure S9. Distribution of deleterious mutations in the sorghum genome.** Distribution of deleterious mutations across sorghum chromosomes. The ratio of deleterious (deleterious, non-conserved deleterious and stop mutation) vs non-deleterious (tolerated and synonymous) variants is higher in pericentromeric regions (gray shading) where recombination is partially suppressed. Green line: smoothing line over the population recombination rate ($\rho$) calculated in 100 kb windows; yellow bars: gene density (100 kb windows); red line: ratio of deleterious to non-deleterious variants (500 kb windows), gray shading: pericentromeric areas.

(A) (B)

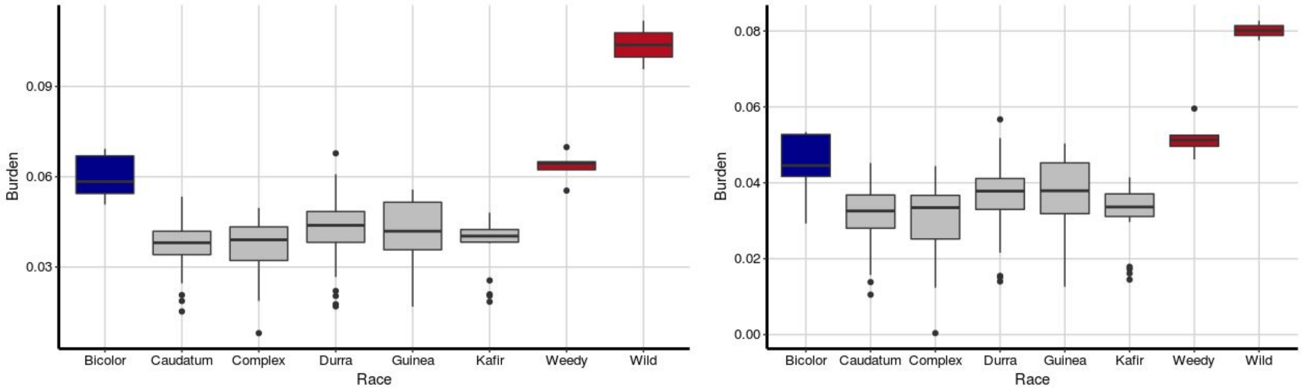


**Figure S10. Deleterious burden in sorghum**. (**A**) Burden was calculated as the number of deleterious alleles ( including stop-codon mutations; either gain or loss, non-conserved deleterious mutations; SIFT <0.05, GERP < 2, and conserved deleterious mutations; Deleterious, SIFT <0.05, GERP > 2), carried by an individual divided by the total number of nonmissing deleterious sites. (**B**) We also calculated burden using only the deleterious category (SIFT < 0.05 and GERP scores > 2).

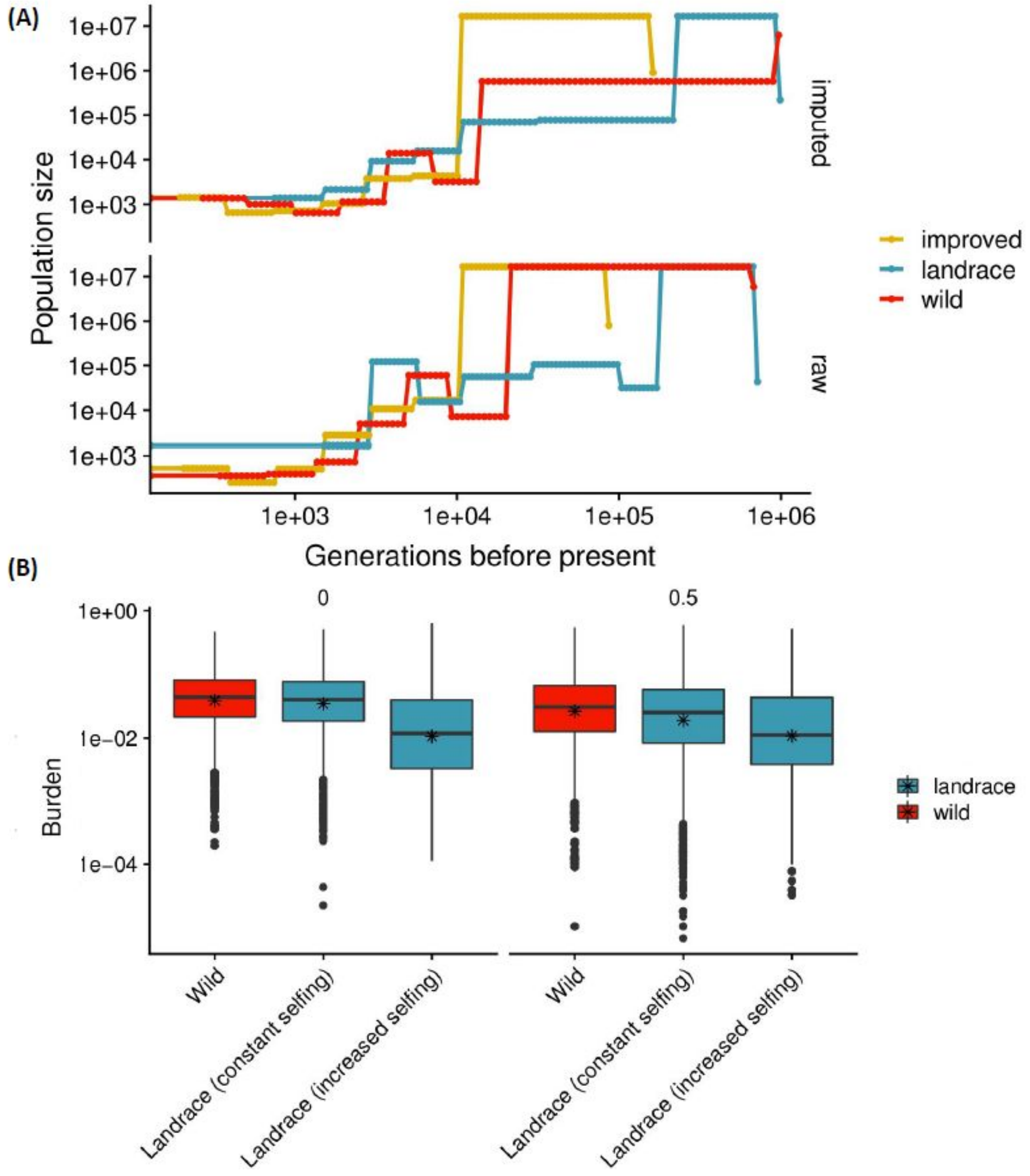


**Figure S11. Sorghum demography**. (**A**) Demographic history of sorghum as inferred by SMC++ [4]. We selected five individuals of each group (wild, landrace, and improved) to ensure a balanced sample size. (**B**) The impact of increased inbreeding on burden during sorghum domestication was explored by changing the inbreeding probability from 0 (0% selfing; 100% outcrossing) to 1 (100% selfing) or from 0.5 (50% selfing) to 1 (100% selfing). The simulations showed that under these scenarios the amount of burden decreased with higher selfing rates.

**(A)**

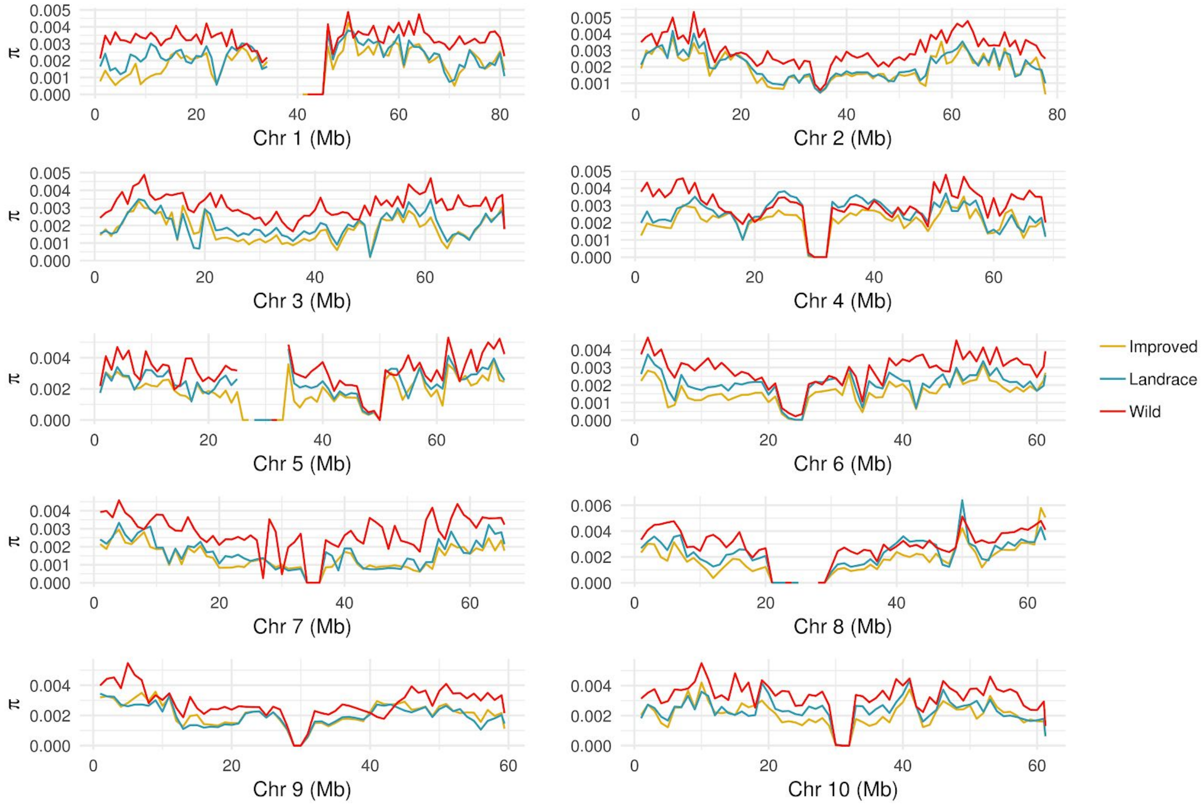


**Figure S12. Nucleotide diversity in sorghum**. (**A**) Nucleotide diversity ( $\pi$ ) was calculated across sorghum chromosomes using 1 Mb bins for improved lines (yellow), landraces (blue), and wild/weedy lines (red). Pi was calculated on a per site basis using VCFtools, we got the sum of nucleotide diversity per window and divided it over the total number of bp that had sequence per group. The denominator was calculated this way to assess the reference bias.

**(A) Average deleterious score**

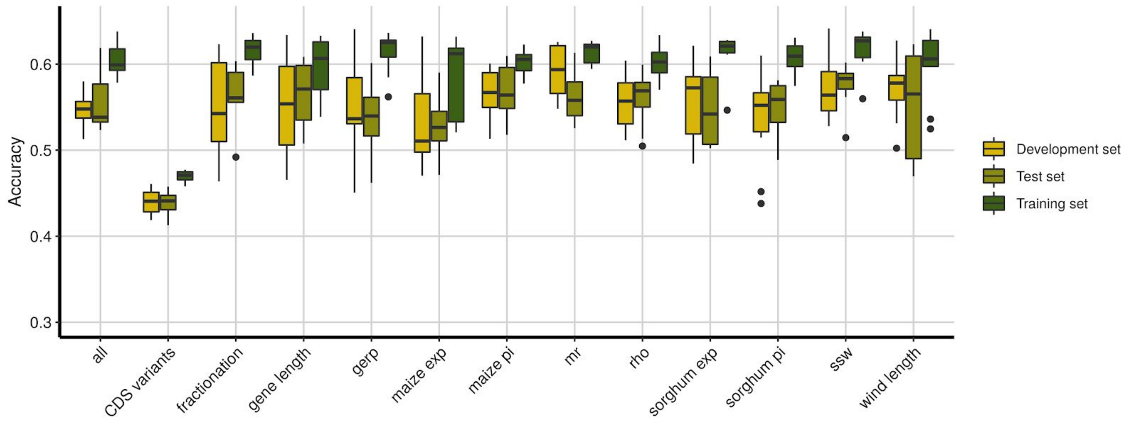


**(B) Syntenic state (Fractionation)**

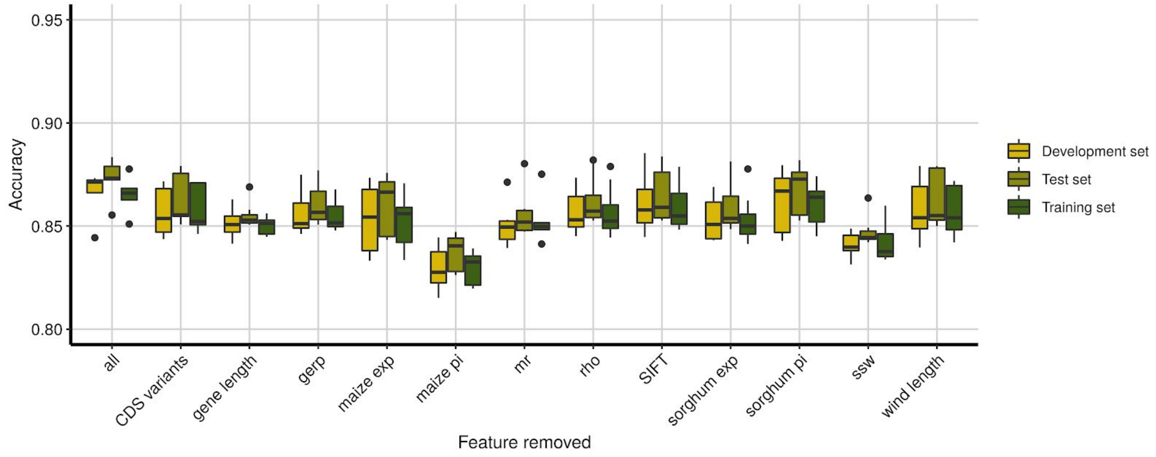


**(C) CNN vs Linear models**

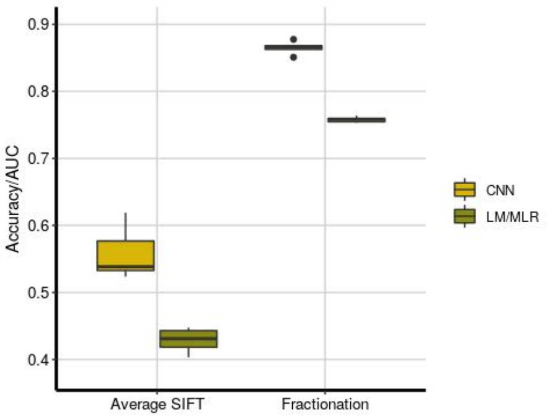


**Fig S13. Convolutional neural network model**. Feature importance when predicting average deleterious score (SIFT) and syntenic state for sorghum. All genomic features are included in the first column (all). For each subsequent column, the feature specified in the x-axis was removed when predicting average deleterious score (**A**) and syntenic state of the focal gene (**B**), respectively. (**C**) Comparison of the prediction accuracies of the CNN models to those from a linear regression (average deleterious score) or multinomial logistic regression models (syntenic state). Accuracy (pearson correlation between predicted and real values) was calculated for average SIFT predictions and area under the curve (AUC) was calculated for syntenic state under a 10-fold cross-validation scheme with 10 replications.

1. Mace, E. S. *et al.* Whole-genome sequencing reveals untapped genetic potential in Africa's indigenous cereal crop sorghum. *Nat. Commun.* **4**, 2320 (2013).
2. Zheng, X. *et al.* A high-performance computing toolset for relatedness and principal component analysis of SNP data. *Bioinformatics* **28**, 3326–3328 (2012).
3. Keightley, P. D. & Jackson, B. C. Inferring the Probability of the Derived vs. the Ancestral Allelic State at a Polymorphic Site. *Genetics* **209**, 897–906 (2018).
4. Terhorst, J., Kamm, J. A. & Song, Y. S. Robust and scalable inference of population history from hundreds of unphased whole genomes. *Nat. Genet.* **49**, 303–309 (2017).
